# The mutation profile of SARS-CoV-2 is primarily shaped by the host antiviral defense

**DOI:** 10.1101/2021.02.02.429486

**Authors:** Cem Azgari, Zeynep Kilinc, Berk Turhan, Defne Circi, Ogun Adebali

## Abstract

Understanding SARS-CoV-2 evolution is a fundamental effort in coping with the COVID-19 pandemic. The virus genomes have been broadly evolving due to the high number of infected hosts world-wide. Mutagenesis and selection are the two inter-dependent mechanisms of virus diversification. However, which mechanisms contribute to the mutation profiles of SARS-CoV-2 remain under-explored. Here, we delineate the contribution of mutagenesis and selection to the genome diversity of SARS-CoV-2 isolates. We generated a comprehensive phylogenetic tree with representative genomes. Instead of counting mutations relative to the reference genome, we identified each mutation event at the nodes of the phylogenetic tree. With this approach, we obtained the mutation events that are independent of each other and generated the mutation profile of SARS-CoV-2 genomes. The results suggest that the heterogeneous mutation patterns are mainly reflections of host (i) antiviral mechanisms that are achieved through APOBEC, ADAR, and ZAP proteins and (ii) probable adaptation against reactive oxygen species.

**Importance:** SARS-CoV-2 genomes are evolving worldwide. Revealing the evolutionary characteristics of SARS-CoV-2 is essential to understand host-virus interactions. Here, we aim to understand whether mutagenesis or selection is the primary driver of SARS-CoV-2 evolution. This study provides an unbiased computational method for profiling and analyzing independently occurring SARS-CoV-2 mutations. The results point out three host antiviral mechanisms shaping the mutational profile of SARS-CoV-2 through APOBEC, ADAR, and ZAP proteins. Besides, reactive oxygen species might have an impact on the SARS-CoV-2 mutagenesis.

## Introduction

The severe acute respiratory syndrome coronavirus 2 (SARS-CoV-2) has spread worldwide since its emergence in December 2019 (Hu, Guo, Zhou, & Shi, 2020), reportedly infected more than 83 million people with a death toll of 1,831,412 as of 4 January 2021, according to World Health Organization (WHO) (https://covid19.who.int/). Studies have been focused on effective treatment of the disease, mostly by the drug re-purposing approach due to the urgency (Shah, Modi, & Sagar, 2020) and by finding a vaccine that will stop the spreading of the virus. Though there are dozens of vaccine candidates in clinical development, the evolutionary potential of the virus might affect the efficacy of the immunizations and treatments. Therefore, understanding the genomic features and mutation dynamics of the virus is crucial to interpret its evolutionary patterns and its response to the available treatments and potential vaccines.

Analyzing virus sequence context and mutations has revealed many critical pieces of information about the characteristics of SARS-CoV-2. For example, the origin of SARS-CoV-2 was linked to bats and pangolins using phylogenetic analyses (Li, Yang, & Ren, 2020; Wu et al., 2020; Zhou et al., 2020; Zhu et al., 2020). Through mutational analyses, some genomic variants of the virus were associated with increased transmissibility (Hou et al., 2020; Volz et al., 2020). In addition, we and others studied the spread of the virus in a variety of countries by tracking the mutation events of sequences over time (Adebali et al., 2020; Popa et al., 2020; Volz et al., 2020).

Some studies also analyzed the mutation profiles of SARS-CoV-2 by counting the mutations on sequences (Graudenzi, Maspero, Angaroni, Piazza, & Ramazzotti, 2020; Popa et al., 2020). Care should be taken when counting mutations to create a mutation profile. Considering that virus genomes are evolutionarily linked to each other, counting all the mutations in the sequences with respect to a reference genome creates a mutation bias towards the most abundant or frequently sequenced isolates. In other words, if a mutation occurs in an ancestral genome, it will also be seen in all of its descendants unless it reverts. When the mutations are called relative to a reference genome, variants of a common origin will be counted multiple times, even though they are linked to a single mutation event. To overcome this issue, we created a phylogenetic tree and assigned only nucleotides that differ from the parent node as a mutation.

In this study, we retrieved more than 65,000 SARS-CoV-2 genome sequences from GISAID (Global Initiative on Sharing All Influenza Data) database (Shu & McCauley, 2017) and clustered them based on their sequence identity. For each cluster, we assigned a sequence as the representative of its cluster. We aligned the representatives, and constructed a phylogenetic tree using a maximum likelihood method. Then, we assigned mutations to the nodes of the phylogenetic tree and counted only the mutations on sequences that were newly acquired with respect to its parent node. Finally, we analyzed mutation profiles and sequence diversity of SARS-CoV-2.

## Results

We retrieved 66,938 genomes, which were all the available genomes in the GISAID database as of July 18^th^, 2020 (Shu & McCauley, 2017). Because multiple sequence alignment and building a tree with more than 65,000 genomes was computationally intense, we initially clustered the genomes and assigned a sequence from each cluster as its representative. By the method, we reduced the genome number to 20,089 without losing major sequence information. To assign mutations, we first aligned the 20,089 representative sequences and built a phylogenetic tree with a maximum likelihood method (see Methods). Then, we reconstructed the phylogenetic tree into a time-resolved phylogenetic tree, which infers the ancestral characters by estimating the molecular clock of the viral genomes (Sagulenko, Puller, & Neher, 2018). After constructing the tree, mutations were assigned based on the differences between the sequences and their parent node **(Figure 1)**. This method enabled us to capture all the mutation events without recounting ancestral mutations. Moreover, we could also identify mutations that occurred repeatedly in different lineages, which would not be possible if the mutations were assigned relative to a reference genome.

**Figure 1:**
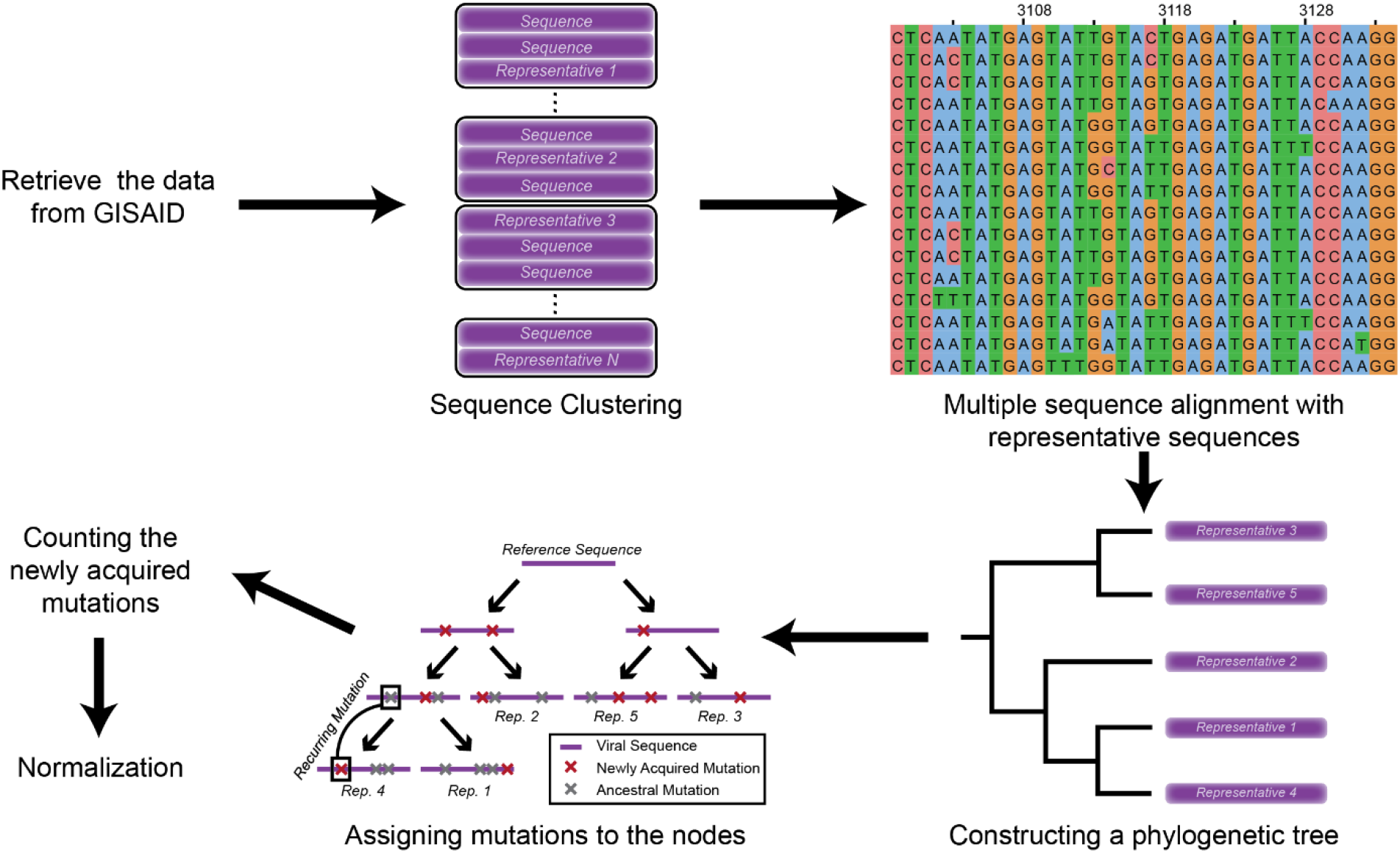
Schematic representation of the methodology.

### Mutation profile of SARS-CoV-2 and potentially related mechanisms

To generate the mutation profile of SARS-CoV-2, we performed mutational signature analysis for all 192 trinucleotide changes using 28,544 mutations from the 35,841 representative sequences and nodes **(Figure 2A)**. We normalized all the trinucleotide changes by the occurrence of the trinucleotide to eliminate any sequence context bias. In general, the most abundant mutational patterns are C>U, G>U, U>C, and A>G substitutions, that are 33.8%, 14.8%, 13.7%, and 9.6% of total substitutions, respectively **(Figure 2A)**.

**Figure 2:**
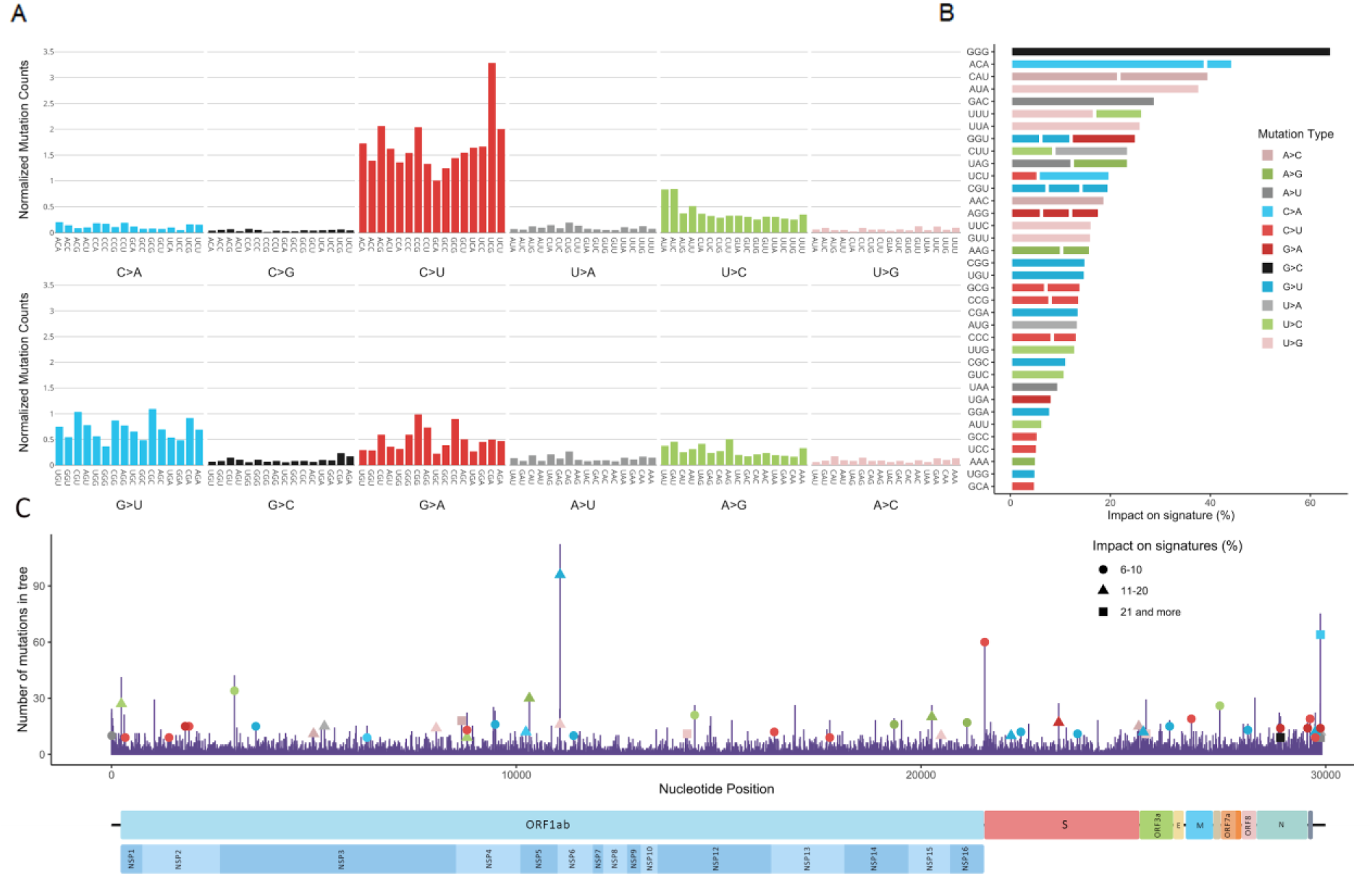
Mutation profile of SARS-CoV-2 genomes. Mutation counts are normalized by the trinucleotide content for each trinucleotide generated from 35,841 (representative sequences and nodes) sequences (A). From unstable positions where more than 8 mutations of the same type at the same position, highly occurred mutations are retrieved. The occurrences of these mutations are divided by the total number of the mutations of the same type and observed in the same trinucleotide. The calculated ratio is used to visualize the impact of highly occurring mutations on signature profile, as percentages (B). Number of mutations observed in the phylogenetic tree, per position. Mutations which have a significant contribution to their signature visualized in part B, are marked according to the mutation count they represent at the position. Marked mutations are colored and reshaped with respect to their mutation type and impact range on their signatures (C).

An enzyme family known for causing C>U substitution is called apolipoprotein B mRNA editing enzyme, catalytic polypeptide-like (APOBEC) family. Enzymes of APOBEC family have an antiviral activity against some RNA viruses including coronaviruses (Harris & Dudley, 2015; Sharma et al., 2015; Woo, Wong, Huang, Lau, & Yuen, 2007). Briefly, they can deaminate cytosine to thymine (uracil in the RNA genome), which can either result in C>U substitution on single-stranded viral RNA (plus strand) or G>A substitution if the mutation occurs on complementary strand (minus strand). In agreement with previous studies, the impact of APOBEC is highly visible at C>U substitutions, while it is relatively low at G>A substitutions (8.8% of total substitutions) (Di Giorgio, Martignano, Torcia, Mattiuz, & Conticello, 2020; Graudenzi et al., 2020; Simmonds, 2020). This result suggests an asymmetric activity for APOBEC enzymes in favor of single-stranded viral RNA. Because virus RNA is frequently present as the plus strand, we see the effect of the APOBEC activity majorly in the form of C>U substitution relative to G>A, which possibly reflects APOBEC activity on the negative strand during RNA replication. Moreover, APOBEC proteins show target inclination towards 5’-[T/U]C-3’ and 5’–CC–3’ motifs while deamination of cytosine (McDaniel et al., 2020). Target sequence preferences of APOBEC proteins are observed in our mutational profile, where the 5 out of 7 highest normalized mutational counts on C>U distributes along 5’-UC-3’ and 5’–CC–3’ motifs **(Figure 2A)**. It is also experimentally found that A1CF RNA editing cofactor, which is APOBEC1 complementation factor, is among the SARS-CoV-2 RNA binders (Schmidt et al., 2020) that strengthens the hypothesis of APOBEC proteins’ activity on the C>U substitutions.

The second most prevalent substitution is G>U, which might be associated with reactive oxygen species (ROS) in APOBEC-related manner. A recent study revealed that DNA damage response mediated by APOBEC3A (a member of APOBEC family) results in ROS production (Niocel, Appourchaux, Nguyen, Delpeuch, & Cimarelli, 2019). ROS can induce oxidative DNA damage, usually transforming guanine into 7,8-dihydro-8-oxo-20-deoxyguanine (oxoguanine), which can pair with adenine and lead to G>U substitution (Molteni, Principi, & Esposito, 2014; Waris & Ahsan, 2006). However, to date, there is no direct evidence of ROS-caused damage in the SARS-CoV-2 genome.

Another mechanism that can mutate the viral genome is adenosine deaminase acting on RNA (ADAR), which is an enzyme that mediates deamination of adenine to inosine (A>I) and later changes to guanine (A>G) (Di Giorgio et al., 2020). A>G (plus strand) and U>C (minus strand) substitutions are observed at similar levels (13.8%, and 9.6% of total substitutions, respectively.) **(Figure 2A)**. ADAR targets dsRNA, and therefore, equivalent levels of ADAR activity is expected to present at both strands (Di Giorgio et al., 2020). The symmetric mutation profile for this pattern strongly suggests that ADAR working on replication RNA is effective in A>G and U>C substitutions.

In the context of trinucleotides, mutations dominantly occurred in U(C>U)G, A(C>U)G, C(C>U)G, U(C>U)U, and A(C>U)A **(Figure 2A)**. Notably, 3 out of the 5 most frequently changed trinucleotides contain CG at their second and third positions. To examine whether these mutations were predominantly located at a single position in the viral genome or are distributed throughout the genome, we identified dynamic positions, where more than 8 recurring mutations were observed. Afterwards, we investigated the contribution of these trinucleotide positions to the mutation profile **(Figure 2B**,**C)**. With some exceptions, most mutations in dynamic positions do not dominate the overall mutation profile. One of the exceptions is G(G>C)G mutations occurred at position 28,883, which correspond to the 64.3% of all mutations occurring on GGG. Although the percentage is high, the number of mutations occurring on GGG trinucleotides is only 14. Similarly G(A>U)C mutations at position 29,869 correspond to the 29% of all mutations occurring on GAC trinucleotide, but the number of mutations of GAC is as low as 31. However, when trinucleotides with a total mutation number exceeding 200 are considered, the position with the highest mutation becomes position 11,083 with U(G>U)U mutations, composing 15% of total mutations of UGU. In conclusion, the mutation profile is not dominated by the switching positions; a position bias on mutation distribution is only observable when the number of mutations of a trinucleotide is low. Several mutations labeled as impactful on signatures **(Figure 2B)** have been investigated for their possible effect on the severity and transmissibility of the virus by various studies. The highest mutational position 11083, has been associated with the severity of the virus (Toyoshima, Nemoto, Matsumoto, Nakamura, & Kiyotani, 2020). Together with 11083, mutations at 23403, 21575, 28881, 28883 positions (Berrio, Gartner, & Wray, 2020; Korber et al., 2020; van Dorp et al., 2020; Yin, 2020) have been associated with significant indication towards selection. Particularly the D614G mutation on Spike protein is associated with the fitness of the virus both by computational and clinical studies (Korber et al., 2020; Plante et al., 2020; Yin, 2020).

### Codon usage of SARS-CoV-2 differentiates in favor of A and U containing codons

Next, we investigated the impact of a potential contribution of codon bias selection on the mutation profile. First, we counted all the formed and altered codons, which we referred to as “form” and “deform”, respectively. We calculated the relative difference between form and deform values for each codon to test potential convergence to host codon usage through mutations **(Figure 3A)**. While UUU, AUA, AUU, and UAU are the intensively formed codons, CCA, UGG, GCU, and ACA are the most diminished ones. These results indicate a dominant forming of A and U containing trinucleotides, whereas G and C containing trinucleotides tend to disappear and reduce in number. In addition, all the codons that are translated into alanine (A) and proline (P) tend to diminish, resulting in lower translation of these amino acids in viral proteins. Considering that all the codons of A and P contain GC and CC in the first and second position, respectively, the reduction in these amino acids is probably related to selection against G and C presence **(Figure 3A)**.

**Figure 3:**
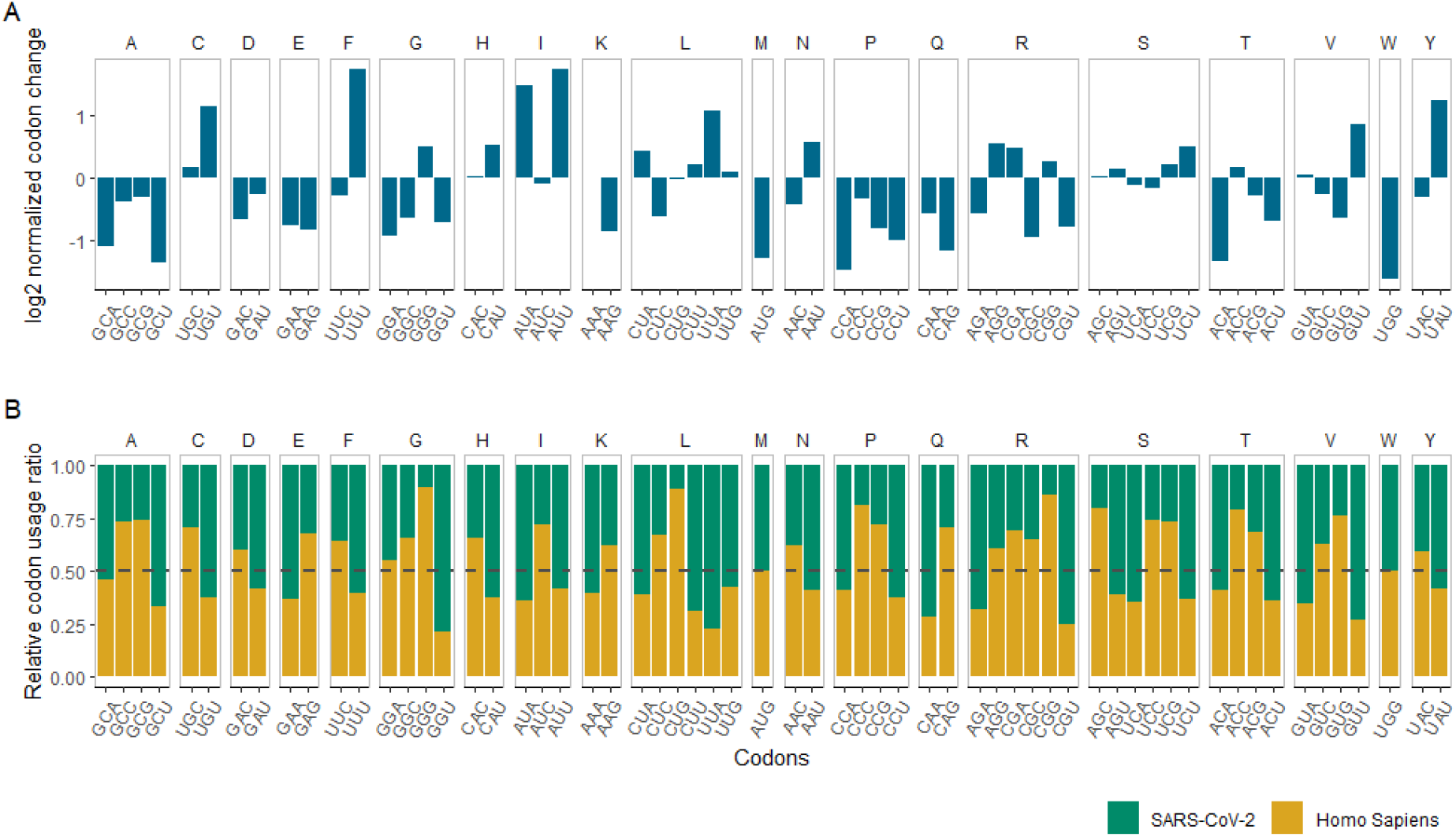
Comparison of codon variations in SARS-CoV-2 phylogenetic tree, human genome and SARS-CoV-2reference genome. (A) Codon variations from mutations in the phylogenetic tree, represented by the ratio of formations over deformations per codon in log2 formation. (B) Relative codon usage percentage between Human and SARS-CoV-2 reference genome.

Human coronaviruses are known to have low GC content (GC%) and SARS-CoV-2 is not an exception with ∼38 GC% (Berkhout & van Hemert, 2015; Xia, 2020). Moreover, it was suggested that the reduction in GC% is an adaptation strategy of SARS-CoV-2, especially to the human lung expressed genes (Y. Li et al., 2020). To determine whether the mutations of SARS-CoV-2 is an adaptation strategy to increase its viability inside the host or just the byproduct of host immune response to the viral RNA, we obtained the human codon usage values (Nakamura, Gojobori, & Ikemura, 2000) and calculated the codon usage of SARS-CoV-2 (see methods). We calculated the relative ratio of these values and grouped codons that are translating the same amino acid **(Figure 3B)**. If the viral genome is to adapt to the host genome, one can hypothesize that the codons that are used dominantly in the host relative to the virus should be formed in the viral genome to increase the similarity, while the codons that are used dominantly in the virus should diminish. GCU, GAA, GGU, CGU codons that are used relatively high in SARS-CoV-2, have the tendency to deform, in agreement with the hypothesis. However, UGU, AUA, UUA, GUU codons that are also used relatively high in SARS-CoV-2, have the tendency to be formed. A similar contradiction is also observed in the codons that are highly used in the human genome. In general, adaptation to the host codon usage cannot explain the formation tendency of the codons. The main driver of the formation tendency is likely to be selection pressure against GC% and thus, A and U increase.

### CG nucleotide deforms, while UU nucleotide forms

After observing higher mutations in trinucleotides that contain CG at their second and third positions, and higher deformation in G and C containing codons, we decided to examine the deformation **(Figure 4A)** and formation **(Figure 4B)** of dinucleotides. Because deformation of a dinucleotide is dependent on its occurrence in the genome, we normalized the deformed value of each dinucleotide with respect to its occurrence in the reference genome. Also suggested by others (Wang et al., 2020; Xia, 2020), CG dinucleotide is the most deformed among all **(Figure 4A)**. Xia et al. attributed the reduction in CG dinucleotide to a protein called zinc finger antiviral protein (ZAP), which binds and mediates the degradation of the viral genome (Xia, 2020). Moreover, the study indicates that SARS-CoV-2 is the most CG deficient betacoronavirus (Xia, 2020). Thus, high CG deformation might be an adaptation of SARS-CoV-2 to escape ZAP under high purifying selection. In addition, UU dinucleotide is formed more than all other dinucleotides. In general, A and U containing dinucleotides are formed, meanwhile C and G containing dinucleotides are deformed.

**Figure 4:**
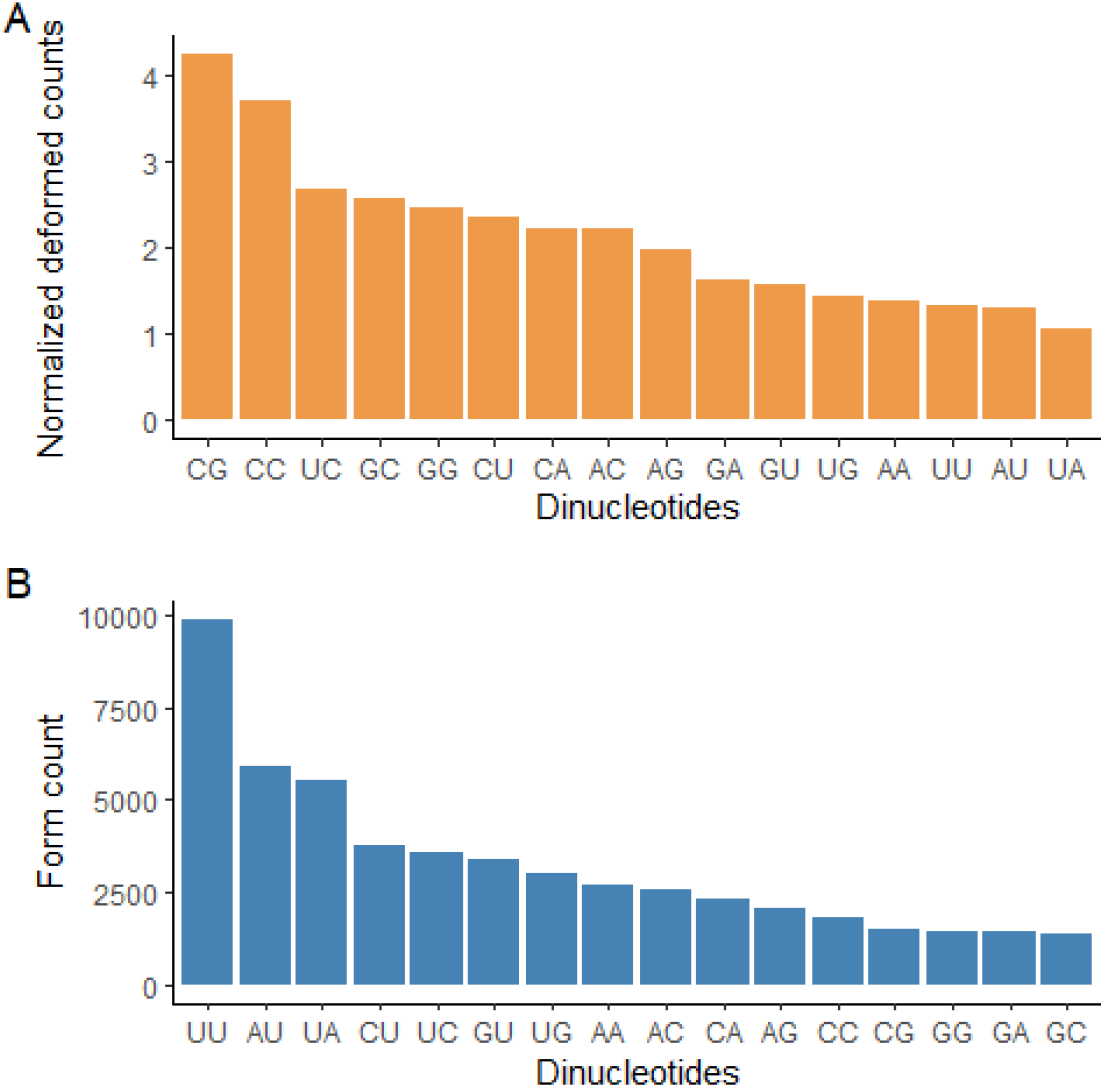
Comparison of dinucleotide formations and deformations retrieved from phylogenetic trees. Deformation ratio of dinucleotides is represented as the ratio of deformation count in the tree over dinucleotide’s abundance in the reference genome(A). As a result of the mutations the relative dinucleotides are formed (B).

## Discussion

The COVID-19 pandemic has been spreading aggressively, killing thousands of people and affecting the daily lives of much more. Moreover, the evolutionary behavior of SARS-CoV-2 might potentially weaken the efficiency of the current treatments and vaccines. Here, we performed a phylogenetic tree-based mutational analysis to assess the contribution of mutagenesis and selection mechanism to SARS-CoV-2 mutation profiles.

The mutation profile of SARS-CoV-2 revealed that C>U, G>U, U>C, and A>G are the dominant substitutions. Based on these mutational patterns, we compiled some potential mechanisms that might be influencing the SARS-CoV-2 viral genome **(Figure 5)**, which are namely APOBEC, ADAR, and ZAP. These mechanisms were linked to SARS-CoV-2 mutagenesis in previous studies as well (Graudenzi et al., 2020; Kosuge, Furusawa-Nishii, Ito, Saito, & Ogasawara, 2020; Xia, 2020). In addition, we suspect that ROS might be a driver of G>U substitutions, however, more studies should be conducted to link ROS to SARS-CoV-2.

**Figure 5:**
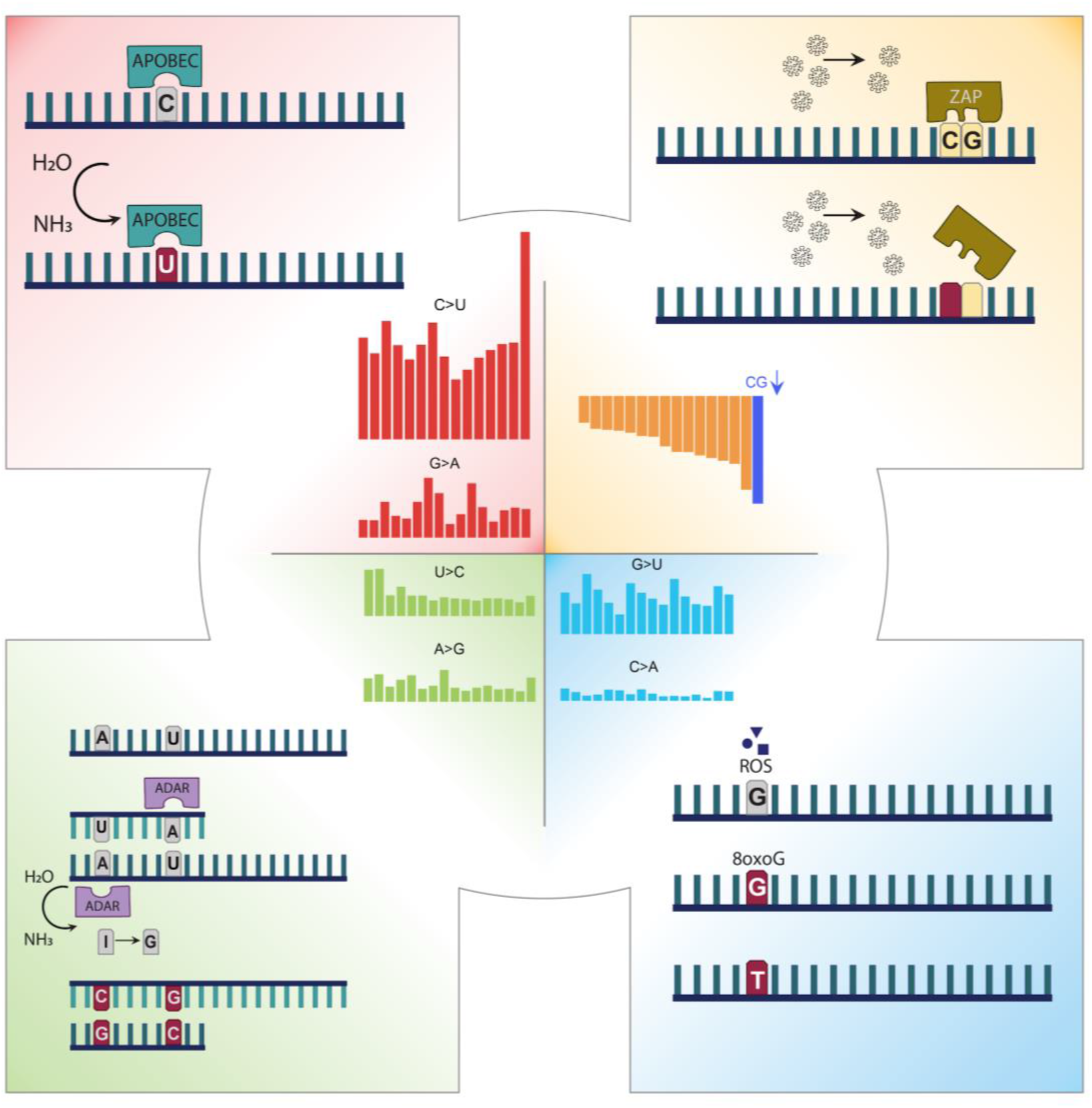
Mechanisms that can alter the sequence context of SARS-CoV-2.

Another aim of this study was to examine the main driver of the mutational patterns of SARS-CoV-2; whether the viral genome is inclined to converge into the host genome or the mechanisms we have discussed are the only contributors to the mutational patterns. Analyses on formed and deformed codons exhibit an increase of A and U and a decrease of G and C containing codons. Furthermore, the comparison between human and SARS-CoV-2 codon usage does not reveal a strong correlation between codon usage percentages and SARS-CoV-2 formation tendency. These results combined, suggest that SARS-CoV-2 genome diverges through RNA editing mechanisms of the host, independently of any adaptive mechanism to increase its genomic similarity to the host genome, which was suggested in another study as well (Y. Li et al., 2020). Then, we examined the formation tendency of dinucleotides. In general, we observed a decrease of G and C and an increase of A and U containing dinucleotides. Strikingly, the deform rate of CG dinucleotides and formation of UU dinucleotides is extremely high. This phenomenon, which was observed in most human viruses (Caudill et al., 2020), was previously associated with the reduction of the hydrogen bonds between strands to achieve more efficient gene expression (Wang et al., 2020).

In conclusion, the mutational profile we generated supported the potential biological mechanisms contributing to the genome diversity of SARS-CoV-2 genomes. Strand asymmetry of some mutation signatures suggested the mechanism acting on the plus RNA strand only. Strand-wise equivalent mutation signature attributed to ADAR is in agreement with its mechanism of action where RNA is affected in the double-strand form. Antiviral responses and selection cannot be distinguished from each other. Host responses against the virus cause mutations in one hand and the reduced targets in the virus genome make it less susceptible to the same antiviral attacks. Although we don’t suggest a direct antiviral mechanism to reduce CG content, the reduced CG content can be explained by the adaptation to the host antiviral mechanism by ZAP. So far, the virus has been affected by the host antiviral mechanisms. Although there are several Spike protein amino acid substitutions that are likely to provide a selection advantage (Chand et al., 2020; Volz et al., 2020), selection hasn’t been a major driving force that can be applied to the entire genome. In the coming months, with a wide administration of the vaccines, it might be possible to see the effect of the vaccination and selection pressure by observing amino acid changes providing an advantage in escaping from immunized hosts.

## Methods

### Data retrieval and mutation assignment

66,938 SARS-CoV-2 genomes and their metadata in the GISAID database, which was dated until 18 July 2020 were retrieved (Shu & McCauley, 2017). Initially, all the sequences with incomplete information (proper date or location of sample collection) in the metadata file were filtered out with a custom python script. Then, the cd-hit program was used to cluster sequences and choose representatives (-c 0.999 -M 0 -T 80) (Fu, Niu, Zhu, Wu, & Li, 2012). 20,089 clusters were created, of which 18,334 contained only a single sequence, while 1,755 of them contained multiple sequences (up to 14,867sequences in a cluster). Then, the first sequence of each cluster was chosen as the representative of that cluster. After representative selection, representative sequences were aligned with the MAFFT algorithm using Augur toolkit (Hadfield et al., 2018; Katoh & Standley, 2016). Wuhan-Hu-1 genome (GenBank: NC_045512.2) was chosen as the reference genome for the alignment. Then, a phylogenetic tree was constructed using IQ-TREE (-fast -n AUTO -m GTR).

The tree then reconstructed into a time-resolved tree using the treetime option of Augur (Hadfield et al., 2018). The sample with the earliest collection date among the representative sequences was chosen as the root and marginal maximum likelihood estimation was used for date inference. The clock rate was applied across the genome to estimate the evolution rate and set to 0.0008, with a standard deviation of 0.0004 and using the date confidence flag to take the uncertainty of divergence time estimates into account. A constant coalescent model was chosen and ‘covariance-aware’ mode of Augur was turned off with no covariance flag. With these parameters, the number of representative sequences reduced to 19,045.

To assign the mutations to the nodes of the time-resolved tree, the ancestral option of Augur, which infers the ancestral sequences, was used by giving the time-resolved tree and the multiple sequence alignment of representative sequences as input (--inference joint).

### Mutation profile analysis

Mutation list was obtained from the phylogenetic tree and includes the mutations observed in each step of the tree. Then, this mutation list was divided into 192 groups based on their 12 mutation types (ie. A>U, G>C) and 16 different trinucleotide content where the mutation is centered (ie. A>U:UAA, G>C:AGU). Each of the 192 mutation groups were normalized with their corresponding trinucleotide count. Trinucleotides were normalized by the number of occurrences in the reference genome. Finally, these normalized mutation count values were plotted within the ggplot2 package (Wickham, 2016) using R language, colored by their corresponding mutation type and trinucleotide content.

Observed mutations are first grouped by their position, mutation type and trinucleotide and frequently observed mutations (more than 8 times) which occur on the same position on the same trinucleotide, are recorded. Then observed mutations are grouped by their mutation type and trinucleotide which resulted in a 192 group as indicated in the mutation profile. By dividing the occurrence counts of frequently observed mutations to their corresponding mutation and trinucleotide groups total occurrence count, the contribution of each mutation to its profile is calculated and the ones who contributed more than 5% are plotted by ggplot2 in R studio.

The number of mutations observed by position is plotted and previously recorded frequently observed mutations **(Figure 2B)** are highlighted with respect to their mutation type and contribution range.

### Counting codon changes and codon usage

By using ancestral mutations from the time-resolved tree, the mutated codons were counted and labeled as deformed count for each codon, while the count of the resulting codons were referred to as formed codons. For each codon type, the ratio between formed and deformed count was taken and plotted in log2 scale by using the ggplot2 package (Wickham, 2016).

Human codon usage table was retrieved (Nakamura et al., 2000), SARS-CoV-2 codon usage table was calculated by a custom R script. Number of occurrences in the reference genome was retrieved for each codon, then, they were grouped by their corresponding amino acids. The ratio of use per codon was calculated by dividing the occurrence of that codon to the sum of itself and its synonymous codons occurrence. Afterwards, the relative ratio of codon usage between Homo Sapiens and SARS-CoV-2 was calculated by dividing the ratio of a codon in one species to the sum of ratio in both species.

### Dinucleotide changes

Observed mutations in the time-resolved tree were used to calculate the number of deformations observed for each dinucleotide. Dinucleotides were formed by these mutations, which were also calculated and recorded as the number of formations for that dinucleotide. Then, observed counts in the reference genome per codon were retrieved. The deformation counts were normalized by their division with their observation counts in the reference genome and plotted with their formation counts by ggplot2 in R studio (Wickham, 2016).

## Code availability

All the codes and the processed data are publicly available at https://github.com/CompGenomeLab/covid19.

## Acknowledgments

This study was supported by EMBO Installation Grant which is funded by TUBITAK. We thank Dr. Stuart James Lucas for his helpful feedback.

## References

Adebali, O., Bircan, A., Circi, D., Islek, B., Kilinc, Z., Selcuk, B., & Turhan, B. (2020). Phylogenetic analysis of SARS-CoV-2 genomes in Turkey. Turk J Biol, 44(3), 146–156. doi:10.3906/biy-2005-35

Berkhout, B., & van Hemert, F. (2015). On the biased nucleotide composition of the human coronavirus RNA genome. Virus Res, 202, 41–47. doi:10.1016/j.virusres.2014.11.031

Berrio, A., Gartner, V., & Wray, G. A. (2020). Positive selection within the genomes of SARS-CoV-2 and other Coronaviruses independent of impact on protein function. PeerJ, 8, e10234.

Caudill, V. R., Qin, S., Winstead, R., Kaur, J., Tisthammer, K., Pineda, E. G., … Pennings, P. S. (2020). CpG-creating mutations are costly in many human viruses. Evol Ecol, 34(3), 339–359. doi:10.1007/s10682-020-10039-z

Chand, M., Hopkins, S., Dabrera, G., Achison, C., Barclay, W., Ferguson, N., … Barrett, J. (2020). Investigation of novel SARS-COV-2 variant: Variant of Concern 202012/01. Retrieved from London: https://assets.publishing.service.gov.uk/government/uploads/system/uploads/attachment_data/file/947048/Technical_Briefing_VOC_SH_NJL2_SH2.pdf

Di Giorgio, S., Martignano, F., Torcia, M. G., Mattiuz, G., & Conticello, S. G. (2020). Evidence for host-dependent RNA editing in the transcriptome of SARS-CoV-2. Sci Adv, 6(25), eabb5813. doi:10.1126/sciadv.abb5813

Fu, L., Niu, B., Zhu, Z., Wu, S., & Li, W. (2012). CD-HIT: accelerated for clustering the next-generation sequencing data. Bioinformatics, 28(23), 3150–3152. doi:10.1093/bioinformatics/bts565

Graudenzi, A., Maspero, D., Angaroni, F., Piazza, R., & Ramazzotti, D. (2020). Mutational signatures and heterogeneous host response revealed via large-scale characterization of SARS-CoV-2 genomic diversity. BioRxiv.

Hadfield, J., Megill, C., Bell, S. M., Huddleston, J., Potter, B., Callender, C., … Neher, R. A. (2018). Nextstrain: real-time tracking of pathogen evolution. Bioinformatics, 34(23), 4121–4123. doi:10.1093/bioinformatics/bty407

Harris, R. S., & Dudley, J. P. (2015). APOBECs and virus restriction. Virology, 479-480, 131–145. doi:10.1016/j.virol.2015.03.012

Hou, Y. J., Chiba, S., Halfmann, P., Ehre, C., Kuroda, M., Dinnon, K. H., 3rd, … Baric, R. S. (2020). SARS-CoV-2 D614G variant exhibits efficient replication ex vivo and transmission in vivo. Science, 370(6523), 1464–1468. doi:10.1126/science.abe8499

Hu, B., Guo, H., Zhou, P., & Shi, Z. L. (2020). Characteristics of SARS-CoV-2 and COVID-19. Nat Rev Microbiol. doi:10.1038/s41579-020-00459-7

Katoh, K., & Standley, D. M. (2016). A simple method to control over-alignment in the MAFFT multiple sequence alignment program. Bioinformatics, 32(13), 1933–1942. doi:10.1093/bioinformatics/btw108

Korber, B., Fischer, W. M., Gnanakaran, S., Yoon, H., Theiler, J., Abfalterer, W., … Montefiori, D. C. (2020). Tracking Changes in SARS-CoV-2 Spike: Evidence that D614G Increases Infectivity of the COVID-19 Virus. Cell, 182(4), 812–827 e819. doi:10.1016/j.cell.2020.06.043

Kosuge, M., Furusawa-Nishii, E., Ito, K., Saito, Y., & Ogasawara, K. (2020). Point mutation bias in SARS-CoV-2 variants results in increased ability to stimulate inflammatory responses. Sci Rep, 10(1), 17766. doi:10.1038/s41598-020-74843-x

Li, C., Yang, Y., & Ren, L. (2020). Genetic evolution analysis of 2019 novel coronavirus and coronavirus from other species. Infect Genet Evol, 82, 104285. doi:10.1016/j.meegid.2020.104285

Li, Y., Yang, X., Wang, N., Wang, H., Yin, B., Yang, X., & Jiang, W. (2020). GC usage of SARS-CoV-2 genes might adapt to the environment of human lung expressed genes. Mol Genet Genomics, 295(6), 1537–1546. doi:10.1007/s00438-020-01719-0

McDaniel, Y. Z., Wang, D., Love, R. P., Adolph, M. B., Mohammadzadeh, N., Chelico, L., & Mansky, L. M. (2020). Deamination hotspots among APOBEC3 family members are defined by both target site sequence context and ssDNA secondary structure. Nucleic acids research, 48(3), 1353–1371.

Molteni, C. G., Principi, N., & Esposito, S. (2014). Reactive oxygen and nitrogen species during viral infections. Free Radic Res, 48(10), 1163–1169. doi:10.3109/10715762.2014.945443

Nakamura, Y., Gojobori, T., & Ikemura, T. (2000). Codon usage tabulated from international DNA sequence databases: status for the year 2000. Nucleic Acids Res, 28(1), 292. doi:10.1093/nar/28.1.292

Niocel, M., Appourchaux, R., Nguyen, X. N., Delpeuch, M., & Cimarelli, A. (2019). The DNA damage induced by the Cytosine Deaminase APOBEC3A Leads to the production of ROS. Sci Rep, 9(1), 4714. doi:10.1038/s41598-019-40941-8

Plante, J. A., Liu, Y., Liu, J., Xia, H., Johnson, B. A., Lokugamage, K. G., … Shi, P. Y. (2020). Spike mutation D614G alters SARS-CoV-2 fitness. Nature. doi:10.1038/s41586-020-2895-3

Popa, A., Genger, J. W., Nicholson, M. D., Penz, T., Schmid, D., Aberle, S. W., … Bergthaler, A. (2020). Genomic epidemiology of superspreading events in Austria reveals mutational dynamics and transmission properties of SARS-CoV-2. Sci Transl Med, 12(573). doi:10.1126/scitranslmed.abe2555

Sagulenko, P., Puller, V., & Neher, R. A. (2018). TreeTime: Maximum-likelihood phylodynamic analysis. Virus Evol, 4(1), vex042. doi:10.1093/ve/vex042

Schmidt, N., Lareau, C. A., Keshishian, H., Ganskih, S., Schneider, C., Hennig, T., … Munschauer, M. (2020). The SARS-CoV-2 RNA-protein interactome in infected human cells. Nat Microbiol. doi:10.1038/s41564-020-00846-z

Shah, B., Modi, P., & Sagar, S. R. (2020). In silico studies on therapeutic agents for COVID-19: Drug repurposing approach. Life Sci, 252, 117652. doi:10.1016/j.lfs.2020.117652

Sharma, S., Patnaik, S. K., Taggart, R. T., Kannisto, E. D., Enriquez, S. M., Gollnick, P., & Baysal, B. E. (2015). APOBEC3A cytidine deaminase induces RNA editing in monocytes and macrophages. Nat Commun, 6, 6881. doi:10.1038/ncomms7881

Shu, Y., & McCauley, J. (2017). GISAID: Global initiative on sharing all influenza data -from vision to reality. Euro Surveill, 22(13). doi:10.2807/1560-7917.ES.2017.22.13.30494

Simmonds, P. (2020). Rampant C-->U Hypermutation in the Genomes of SARS-CoV-2 and Other Coronaviruses: Causes and Consequences for Their Short- and Long-Term Evolutionary Trajectories. mSphere, 5(3). doi:10.1128/mSphere.00408-20

Toyoshima, Y., Nemoto, K., Matsumoto, S., Nakamura, Y., & Kiyotani, K. (2020). SARS-CoV-2 genomic variations associated with mortality rate of COVID-19. J Hum Genet, 65(12), 1075–1082. doi:10.1038/s10038-020-0808-9

van Dorp, L., Acman, M., Richard, D., Shaw, L. P., Ford, C. E., Ormond, L., … Balloux, F. (2020). Emergence of genomic diversity and recurrent mutations in SARS-CoV-2. Infect Genet Evol, 83, 104351. doi:10.1016/j.meegid.2020.104351

Volz, E., Hill, V., McCrone, J. T., Price, A., Jorgensen, D., O’Toole, A., … Connor, T. R. (2020). Evaluating the Effects of SARS-CoV-2 Spike Mutation D614G on Transmissibility and Pathogenicity. Cell. doi:10.1016/j.cell.2020.11.020

Wang, Y., Mao, J.-M., Wang, G.-D., Qiu, Z., Yao, Q., & Chen, K.-P. (2020). Human SARS-CoV-2 has evolved to reduce CG dinucleotide in its open reading frames.

Waris, G., & Ahsan, H. (2006). Reactive oxygen species: role in the development of cancer and various chronic conditions. J Carcinog, 5, 14. doi:10.1186/1477-3163-5-14

Wickham, H. (2016). ggplot2: elegant graphics for data analysis: springer.

Woo, P. C., Wong, B. H., Huang, Y., Lau, S. K., & Yuen, K. Y. (2007). Cytosine deamination and selection of CpG suppressed clones are the two major independent biological forces that shape codon usage bias in coronaviruses. Virology, 369(2), 431–442. doi:10.1016/j.virol.2007.08.010

Wu, F., Zhao, S., Yu, B., Chen, Y. M., Wang, W., Song, Z. G., … Zhang, Y. Z. (2020). Author Correction: A new coronavirus associated with human respiratory disease in China. Nature, 580(7803), E7. doi:10.1038/s41586-020-2202-3

Xia, X. (2020). Extreme Genomic CpG Deficiency in SARS-CoV-2 and Evasion of Host Antiviral Defense. Mol Biol Evol, 37(9), 2699–2705. doi:10.1093/molbev/msaa094

Yin, C. (2020). Genotyping coronavirus SARS-CoV-2: methods and implications. Genomics, 112(5), 3588–3596. doi:10.1016/j.ygeno.2020.04.016

Zhou, P., Yang, X. L., Wang, X. G., Hu, B., Zhang, L., Zhang, W., … Shi, Z. L. (2020). A pneumonia outbreak associated with a new coronavirus of probable bat origin. Nature, 579(7798), 270–273. doi:10.1038/s41586-020-2012-7

Zhu, N., Zhang, D., Wang, W., Li, X., Yang, B., Song, J., … Research, T. (2020). A Novel Coronavirus from Patients with Pneumonia in China, 2019. N Engl J Med, 382(8), 727–733. doi:10.1056/NEJMoa2001017

